# Pulsed blue light and phage therapy: A novel synergistic bactericide

**DOI:** 10.1101/2024.07.29.605651

**Authors:** Amit Rimon, Jonathan Belin, Ortal Yerushalmy, Sivan Alkalay-Oren, Yonatan Eavri, Anatoly Shapochnikov, Shunit Coppenhagen-Glazer, Ronen Hazan, Lilach Gavish

## Abstract

Antibiotic-resistant *Pseudomonas aeruginosa* (PA) is a critical health threat. Novel treatment approaches are urgently required in this post-antibiotic era. In the current study, we investigated the bactericidal combinatorial potential of two non-antibiotic alternative approaches: phage therapy and pulsed blue light (PBL). Bacteriophages (phages), are viruses that specifically infect and lyse bacteria without harming eukaryotic cells. Pulsed blue light (PBL) alters bacterial membranes and was clinically shown to be innocuous to the skin in low doses. Here, using a low dose 457nm, 33KHz PBL combined with specific PA targeting phages, we demonstrated a synergistic effect that achieved complete inhibition of planktonic bacteria and a 40% reduction in formed biofilms. As part of this study, we also developed a user-friendly python-based tool for extraction of growth curve outcomes. *In vivo* studies are warranted for further validation of this combinatorial treatment. This approach may lead to a novel, antibiotic complementary modality to help patients suffering from difficult-to-treat antibiotic-resistant infections.

**One Sentence Summary:** Low-dose pulsed blue light and phage therapy have a synergistic bactericidal effect on *Pseudomonas aeruginosa* planktonic cultures and formed biofilm

## 1 INTRODUCTION

### 1.1 Antibiotic-Resistant Bacteria and Biofilms

The rise in prevalence and propagation of antibiotic-resistant bacteria is a critical health concern, with a direct association of 1.27 million deaths in 2019 alone^1^. The genetic plasticity of bacterial populations, facilitated by selective pressures from antibiotic over-usage in clinical, agricultural, and environmental applications, results in rapid bacterial adaptation and resistance mechanisms ^2–4^. This phenomenon drastically curtails the efficacy of conventional antibiotic therapies particularly in biofilms, structured bacterial communities encased in a self-produced extracellular matrix. These microbial aggregates foster increased resistance to antimicrobial agents and host immune responses^5,6^ and pose substantial challenges in clinical and industrial settings^7–9^. *Pseudomonas aeruginosa (PA),* a gram-negative rod, is one of the 6 leading pathogens for death associated with antibiotic resistance^1^, and appears in a biofilm form, particularly in burns and other wounds ^10^. Thus, novel treatments, approaches, or combinations of approaches are needed at this time against it. In this study, we focus on two such approaches and demonstrate their synergism against PA.

### 1.2 Phage Therapy

A rising reinvigorated approach against resistant bacteria is the use of bacteriophages (phages ^11^, viruses that kill bacteria without harming eukaryotic cells and which are the most prominent biological entity with estimated 10^31^ particles worldwide^12^. Unlike antibiotics, phages target specific bacterial strains, while largely sparing other microbiota^12^. Among the advantages of using phages as therapeutic agents is their ability to penetrate biofilm structures^13^ and most importantly, bacterial phage resistance can be handled through novel phage isolation, phage engineering, or naturally by evolutionary changes^14^. Phage therapy has proven effective against several antibiotic-resistant pathogens including *Pseudomonas aeruginosa*^15–17^*, Staphylococcus aureus*^18^, and *Cutibacterium acnes*^19–21^, particularly in skin-related infections. Moreover, phage therapy has been successfully used in compassionate cases and case series^15,22–25^ including by our team in the Israeli Phage Therapy Center (IPTC), of non-healing infections caused by antibiotic-resistant bacteria^26,27^.

### 1.3 Pulsed Blue Light (PBL)

Another promising approach to combat resistant bacteria is antimicrobial blue light, that was shown to be effective against both gram- positive and gram-negative bacteria notably against antibiotic resistant PA strains^28–30^. The blue light photons were shown to be absorbed by bacterial chromophores such as porphyrins resulting in excessive reactive oxygen species (ROS) that alter the bacterial membranes^31^. Blue light in low doses was determined clinically to be innocuous to the skin^32^. Pulsed blue light (PBL) was shown to have greater bactericidal efficacy compared to continuous blue light in much lower power intensity and energy doses in planktonic cultures^33^ and was shown to partially disrupt and disassemble biofilm structures^34^.

### 1.4 Study Rationale and Objective

In view of the increase in antibiotic-resistant bacteria and fewer available new antibiotics^35^ coupled with the fact that PBL and phages eradicate bacteria through different mechanisms, we hypothesize that a combination of these interventions may culminate in a synergistic bactericidal effect.

The objective of the current study was to determine the efficacy of this novel approach on planktonic and biofilm PA cultures.

## 2 RESULTS

### 2.1 Selected phages for the study

This study used two phages specific for PA, lytic (PASA16) and lysogenic (PAShipCat1). The phage PASA16 was previously isolated and characterized in our lab^36^ and used in various clinical compassionate treatments^15^. As described previously^36^, PASA16 has a 66,127 bp long genome (GenBank accession number MT933737.1) with a GC content of 55.6% and contains 90 coding DNA sequences (CDS).

PAShipCat1 is a novel lysogenic phage isolated and characterized in this work. This phage has a 50602 bp long genome (GenBank accession number PP067092), with a GC content of 59.3% and 82 coding DNA sequences (CDS). According to its genome sequence, PAShipCat1 belongs to the Caudoviricetes class (Figure 3). Since the genome of PAShipCat1 is found as a prophage in many PA bacteria and encodes a C-like Repressor, it probably has lysogenic abilities^37^.

These two phages were selected deliberately due to their moderate effects, which allowed us to observe synergistic effects with the PBL. The phage PAShipCat1 was used in the planktonic-culture experiment (Figure 4A). In the biofilm experiment, we used PASA16, which had a milder efficacy against biofilm (Figure 4B), although it is highly active against planktonic cultures (Fig 4A).

**Fig. 1.**
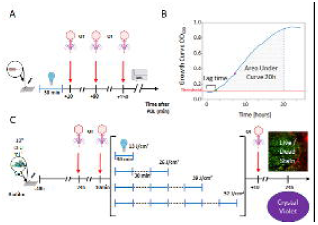
Experimental Design and Outcomes. (A) PAShipCat1 phage was added 10, 90, or 150 minutes after 30 minutes of pulsed blue light (PBL) treatment to planktonic *P. aeruginosa* (PA14). The bacteria were transferred to a plate reader for 24 hours to produce growth curves. (B) Measures of bactericidal effect were extracted from the growth curves including ‘Lag time’ (time to threshold) and ‘Area under the curve at 20 hours (AUC 20). (C) Biofilms of PA14 were formed and PASA16 phage was added either 24 hours or 10 minutes prior to PBL or 10 minutes after. PBL included multiple 30-minute irradiation (blue line) with 30 minute intervals (dotted line). Crystal violet or live-dead stain were used to evaluate the biofilm biomass and viability. This figure was created with Biorender.com

**Fig. 2.**
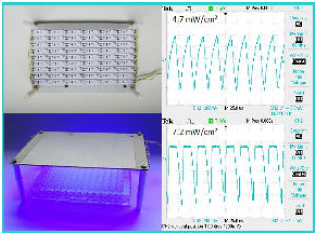
Pulsed Blue Light Source. Photograph of the pulsed blue light source used in this study designed to homogenously irradiate a 96-well plate at a distance of 5 cm within a humid incubator. A total of 153 royal blue LEDs (457nm) were attached to an aluminum surface and electrically interconnected. Power was supplied via an LED driver with a 555 pulse generator module. Waveform and frequency were measured with an oscilloscope connected to a light detector. Pulses were generated at a frequency of 33kHz with a 50% duty cycle and their waveforms were triangular and rectangular at 4.7 and 7.2 mW/cm² respectively.

**Fig. 3:**
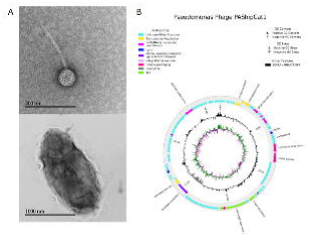
PAShipCat1 Sequence and Structure. The anti-*Pseudomonas aeruginosa* phage, PAShipCat1. (A) TEM imaging reveals a typical structure of Caudoviricetes class-phage with an icosahedral head and a tail, in agreement with the genomic data (B) which also placed it within the Caudoviricetes family taxonomically. Due to its possession of a c-like repressor and phage homologs detected in other bacterial genomes, it has most probably lysogenic characteristics.

**Fig. 4:**
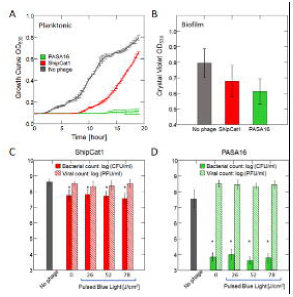
Comparative Phage Bactericidal Efficacy and Stability under PBL. (A) Bactericidal efficacy variation among phages: PASA16 demonstrated potent elimination of PA in planktonic culture, while PAShipCat1 exhibited a moderate effect. (B) In a biofilm setting, PASA16 outperformed PAShipCat1, showcasing higher efficiency. (C-D) Effect of Pulsed Blue Light (PBL) on Phage Stability: Viable counts of planktonic *P. aeruginosa* infected with PAShipCat1 or PASA16 phages pre-exposed to 7.2 mW/cm² pulsed blue light for up to 3 hours (78 J/cm²). Bar graphs represent mean ± SD, n=5 biological repeats per dose (average of triplicates). Note that the phages significantly reduced the bacterial count, while PBL did not influence viral count or the bactericidal effect of the phage. *p<0.001 by ANOVA Tukey as post hoc.

### 2.2 PBL has no direct effect on phage number or activity

Initially, we assessed the effect of PBL on the phages by irradiating them followed by quantification of phage titer. The titer of PAShipCat1 or PASA16 phages measured in PFU/ml was not reduced in up to 3 hours of PBL irradiation (p>0.5 by 1-way ANOVA for both phages). The bactericidal activity of these irradiated phages was not impaired as determined by CFU (p<0.001 for all PBL doses) (Figure 4C and D and supplemental Table 1).

### 2.3 Timing of Administration and Dose/Concentration Response

After determining that 457nm 33kHz PBL irradiation does not directly affect phage activity, we proceeded to evaluate PBL/Phage combination treatments. The timing of phage addition to planktonic *P. aeruginosa* (10^6^ CFU) in relation to PBL was tested by adding PAShipCat1 phage (10^6^ PFU) at 10, 90, or 150 minutes post the 30’ PBL treatment (8.5 J/cm^2^) (Figure 1A). While PBL and phage alone, significantly reduced the area under the curve (AUC) at 20h post treatment compared to the controls by 28% and 51% respectively, their combination reduced it by 73% and the best timing was determined at 10-minutes post PBL (Figure 5A and supplemental Table 2).

**Fig. 5:**
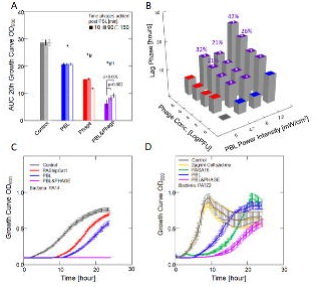
PBL and Phage Synergism is Dependent on the Interval Duration between Interventions, phage concentration, and PBL dose. (A) The bactericidal effect in response to 8.5 J/cm^2^ PBL followed by addition of 10^6^ PFU Phage 10, 90, or 150 minutes later. Bar=mean ± SD area under growth curve (AUC) at 20 hours, n=4 (each biological repeat was an average of triplicate technical repeats). Addition of phage at 10 minutes was the most effective. *, #, † p<0.001 from control, PBL, phage respectively by 2-way ANOVA with Tukey as post-hoc test. (B) The bactericidal effect in response to different combinations of PBL dose × phage concentration: Bacteria were irradiated for 30 minutes with 0, 4.7, 6, or 7.2 mW/cm^2^ PBL (0, 8.5, 10.8, or 13 J/cm^2^) and 10 minutes later, phages were added at 0, 10^2^, 10^4^, 10^6^, or 10^8^ PFU/ml. Bar=mean ± SD lag time [hours] over control. Bactericidal effects were directly associated with dose and concentration. p<0.001 for all comparisons by 2-way ANOVA with Tukey as post hoc. *additive effect (±10% of expected); %Synergistic effect (> 20% over expected). (C-D) Growth curves by OD_600_: PA14 or antibiotic resistant PATZ2 were treated with PAShipCat1 or PASA16 at 10 PFU/ml respectively with or without 30 minutes PBL 7.2 mW/cm^2^ both showing synergistic bactericidal effect. n =5 (each an average of triplicate technical repeats).

Using the above protocol, we tested the effect of different phage concentrations combined with different PBL power intensities on bacterial growth curve lag phase. Phage alone significantly extended the lag phase as did PBL alone (p<0.001 for all concentrations and power intensities). Combining PBL and phages resulted in an additive effect for 10^2^/10^4^ PFU/ml and up to 144% synergistic effect for greater concentrations (Figure 5B) with no bacterial growth observed at the highest dose/concentration combination (Figure 5C). This combination reduced the CFU by 9 logs in PA14 and achieved significant results also when applied to the 2 μg/ml Ceftazidime-resistant PATZ2 (Figure 5D and supplemental Table 3).

### 2.4 Biofilm Viability and Biomass

Next, we assessed the effect of the PBL-phage combination in a more challenging setting, formed biofilm. Initially, we tested the timing of phage addition in relation to the PBL doses (Figure 1C). Using crystal violet stain, we found that both PBL and phage reduced biofilm biomass with the most efficient protocol being phage added immediately before PBL regardless of the dose (p<0.001 across doses, presence of phage, and timing of phage addition by 3-way ANOVA) (Figure 6A). Based on these results we chose to add the phage immediately before a 26J/cm^2^ PBL treatment for the next stage. Using live/dead stain we found that all treatments significantly decreased the biofilm viability (p<0.001 vs controls). While phage alone decreased biofilm viability by 50% of control, addition of PBL further reduced the biofilm viability to 19% (p=0.012 PBL&phage vs phage alone) (Figure 6B and supplemental Table 4).

**Fig. 6:**
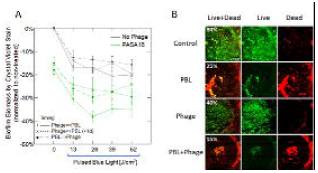
Biofilm Viability and Biomass is Effectively Reduced Following PBL and Phage Treatment. (A) Graph showing the reduction in biofilm biomass in response to PBL dose immediately before or 1 day after addition of phage by crystal violet staining. Biofilms were irradiated with 0-52 J/cm^2^ PBL (see figure 2C for detailed experimental design). Data points = mean+SD %reduction in crystal violet stain levels (normalized to control). Lines added for visualization. The most effective combination was a PBL dose of 26 J/cm^2^ immediately after phage administration. n=5 biological repeats per dose (each an average of 3 technical repeats). (B) Representative live-dead fluorescently stained biofilm samples from each experimental group visualized at ×4 magnification with %Live of total determined by the method of Robertson et al ^67^. Maximal intensity projection of the Z-stack constructed by confocal microscopy was extracted from 17 sections in 12.5μm intervals. Note that the samples treated by PBL exhibit significantly higher red staining demonstrating a reduction in biofilm viability.

## 3 DISCUSSION

The results of this study show that both planktonic and biofilm cultures of PA are highly susceptible to phage therapy combined with pulsed blue light. The synergistic effect was found to result in 99% reduction of planktonic cultures and 40% of preformed biofilms with the best bactericidal effects obtained at a dose of 26J/cm^2^ with 10^8^ PFU/ml phage added within 10 minutes of PBL administration. This bactericidal combination of PA-targeted phage therapy and non-risk low-dose pulsed blue light, can be used against superficial wound infections such as burns, acne, diabetic foot ulcers, cellulitis and others that pose an escalating challenge due to antibiotic resistance^1,38^. Moreover, this combined approach can provide a novel avenue for further enhancing biofilm management^7,39^.

### 3.1 Phages and antibiotics

Currently, phage therapy is usually combined with antibiotic treatment ^40^. Lin et al. reported that the eradication level is dependent on the PA strain and the antibiotic used. For example, phage PEV20 achieved complete inhibition in planktonic cultures when combined with ciprofloxacin but not with amikacin and colistin^41^. However, Knezevic et al., reported that combination with ciprofloxacin leads to only a 3.3 log reduction in PA^42^. Similarly, in this study we demonstrated complete inhibition of planktonic PA14 but partial inhibition of PATZ2. Interestingly, antibiotics alone were ineffective in reducing *Pseudomonas* biofilm biomass^5,43^. However, Chaudhry et al. demonstrated that combining 10 µg/ml ciprofloxacin with phages resulted in a 47-74% biomass reduction in a clinical cystic fibrosis *P. aeruginosa* strain originated biofilm, higher than the 40% synergistic effect found in the current study^43,44^. Although additional PBL dose did not increase the level of biomass eradication (Figure 6A), increasing the power intensity may result in increased eradication efficacy (Figure 5B) which will be the focus of future studies.

### 3.2 Blue Light

While blue light may be harmful to the eyes, skin exposure was clinically determined to be innocuous^32^. In a clinical study, healthy volunteers were exposed for 5 consecutive days to a daily dose of 100J/cm^2^ of 420 nm blue light. No indication of DNA damage, inflammation, or apoptosis were detected in their skin biopsies, although transient keratinocyte melanogenesis and vacuolization was observed^45^. However, in vitro, blue light at wavelength of 453nm, exerted cytotoxic effects on human keratinocytes in doses higher than 500 J/cm^2^ ^46^– 20 times higher than the doses used in this study. In another cell culture study, blue light of 470nm was reported to slightly decrease cell count and viability in fibroblasts using a power of 100mW/cm^2^ culminating to a dose of 110J/cm^2^ ^47^– 12 times the power and 4 times the dose used here. These studies indicate that the wavelength and energetic parameters used in the current study are not expected to be harmful to the skin. The bactericidal effect of continuous blue light was extensively studied with special attention to the tradeoff between power and dose to avoid a photothermal effect: Rupel et al used 455nm blue light, irradiance of 300mW/cm^2^ and dose of 60 or 120 J/cm^2^ to irradiated planktonic cultures of PA and achieved inhibition for 24h^48^. In the current study we achieved the same by using a similar wavelength, 7.2mW/cm^2^, and dose of 13J/cm^2^. Pulsed light allows using higher peak power densities compared to those that could be used in continuous waves, without exposing the skin to excessive tissue heating^49^. Indeed, PBL in very low average irradiances and doses was determined to be superior to continuous wave. Using multiple 30-minute sessions of 450nm PBL at doses of 7.6 J/cm^2^ or less, Bumah et al reported reduction of 7 log CFU/ml of planktonic *C. acne* and MRSA and determined that PBL was superior to continuous wave in similar doses. Additionally, they reported that PBL suppressed *C.acnes* and MRSA biofilms by 64% and 53% respectively^34,50^. In the current study, addition of phage to PBL in similar wavelength and energetic parameters achieved 40% suppression of pre-formed PA biofilms.

### 3.3 Direct effect of PBL on phages

We found that pulsed blue light did not affect viral count or activity in neither PASA16 or PAShipCat1, even when irradiated for extended durations and high dose (up to 3 hours, 7.2mW/cm^2^, and 39 J/cm^2^). However, previous reports indicated that several single stranded RNA viruses can be inactivated. Enwemeka et al reported that 405-450 nm PBL with 12 mW/cm^2^ (130 J/cm^2^) ^51^ can effectively inactivate strains of coronavirus. Tomb et al and Richardson et al reported that continuous blue light can inactivate feline calicivirus^52^ and murine leukemia virus^53^ respectively. It was found that light did not directly affect the viral envelope or viral genome but rather impaired the reverse transcription complex^53^. Both PASA16 or PAShipCat1 have a double stranded DNA genome, thereby they do not include the RT complex required for multiplication in RNA viruses. This lack of RT complex and the much lower power level and total dose applied, may explain why these phages were not susceptible to pulsed blue light used in the current study.

### 3.4 Practical considerations

We found that the most effective timing is PBL immediately after phage administration in biofilm cultures (Figure 5-6) and that the synergistic effect was still prominent even when phage was administered 24h before PBL such as in the case of intravenous, or oral phage delivery^54,55^. Our study employed clinically applicable pulsed blue light power density and pulse frequency^34,47,56^, making our findings relevant to potential clinical applications. Notably, one of the two phages utilized, PASA16, has been previously involved in various clinical compassionate treatments^15^.

### 3.5 Mechanism

The proposed synergism between phage therapy and pulsed blue light could involve at least four possible mechanisms. Firstly, similar to UV, PBL might induce lysogen activation, triggering dormant phages within bacterial cells^57^, thereby enhancing overall phage activity. Secondly, a significant change in the Multiplicity of Infection (MOI), the amount of phages for each bacterium, could illustrate the association between phage activity and bacterial numbers^58^, with lower initial counts leading to more efficient phage activity. A third potential scenario involves occupying all of the cell’s DNA repair mechanisms, which could impede bacterial recovery and proliferation, limiting the spread of the infection. Finally, membrane instability induced by pulsed blue light^31,56^ might facilitate the activity of phage lytic enzymes, promoting the rapid destruction of bacterial cells. Further studies are warranted to explore which mechanism(s) is responsible to this synergistic effect.

### 3.6 Limitations

Our study’s limitations stem from the exclusive focus on the PA strains PA14 and PATZ2 with only two phages, limiting the generalizability of our findings. Prior research highlighted variable phage effectiveness in reducing *Pseudomonas* cultures or biofilm biomass, emphasizing the specificity of strains and phages ^59,60^. Another limitation is the 30-minute treatment duration in our study, potentially being lengthy. Subsequent experiments should explore shorter durations and higher power settings. Additionally, we limited ourselves to the use of 457nm wavelength although shorter wavelengths were shown to be more effective, to preclude safety concerns which arise with UV wavelengths ^61^. Nevertheless, these results are promising enough to pursue combinations of PBL and phages as a novel therapeutic to superficial infections.

In conclusion, we present a bactericidal novel synergistic combination of phage therapy and low-dose pulsed blue light demonstrating effectiveness in both planktonic and pre-grown biofilm of *Pseudomonas aeruginosa*. Future research should include diverse bacterial strains, additional phages, and a comprehensive exploration of underlying mechanisms. Finally, clinical trials should assess the clinical implications of the treatment suggested herein for patients inflicted with antibiotic resistant superficial infectious conditions.

## 4 MATERIALS AND METHODS

### 4.1 Study Design

In this study we used a combinatorial dose-response design (phage concentration × PBL power density) to achieve a bactericidal effect over *P*. *aeruginosa* planktonic cultures in the logarithmic phase and pre-formed biofilms (Figure 1A-C).

### 4.2 Bacteria

#### 4.2.1 Bacterial strains

The bacteria used here was the reference *Pseudomonas aeruginosa* strain PA14^62^, obtained from our laboratory stock collection and PATZ2 that was isolated from a clinical phage therapy case in our institution^15^ and was determined to be resistant to 2μg/ml Ceftazidime (Panpharma Z.I, Luitre, France).

#### 4.2.2 Planktonic bacteria

Bacteria were grown, from a -80°C stock, overnight in Luria Bertani (LB) medium at 37°C under conditions of continuous agitation, diluted 1:1000, and re-grown in LB. Bacteria grown to mid-logarithmic (OD_600_ ∼= 0.5) phase were used for the experiments.

#### 4.2.3 Biofilm

Bacteria were grown overnight, and 200 μl of 10^8^ CFU/ml per well were transferred into the inner wells of a 96-well plate, and grown for 48h. Outer wells were filled with double distilled water to prevent inhomogeneous evaporation^63^.

### 4.3 Phage isolation and characterization

#### 4.3.1 Phage isolation

Isolation of phages was performed using the standard double-layered agar method^64^. Briefly, an environmental sample was centrifuged (centrifuge 5430R, rotor FA-45-24-11HS; Eppendorf, Hamburg, Germany) at 10,000*g* for 10 min, and the supernatant was filtered through 0.22-μm-pore-size filters (Merck Millipore, Rahway, NJ). Exponentially grown PA14 (10^8^ CFU/ml) cells were inoculated with a filtered sample, incubated for 24 h at 37°C and then filtered. The lysate was diluted in LB broth and spotted on a PA14 bacterial lawn grown on soft 0.6% agarose and incubated overnight at 37°C. Observed plaques were transferred into broth using a sterile loop and stored in LB at 4°C.

#### 4.3.2 Phage sequencing

The phage genome sequencing was performed as previously described^19^. Briefly, the phage DNA isolation kit (Norgen Biotek, Thorold, Canada) and the Illumina Nextera XT DNA kit (Illumina, San Diego, CA) were used to extract DNA and prepare sequence libraries respectively. A flow cell (1 × 150 bp single-end reads) was used for normalization, pooling, and tagging and the Illumina NextSeq 500 platform was used for sequencing. Trimming, quality control, read assembly, and analyses were performed using Geneious Prime® 2023.2.1 and its plugins (https://www.geneious.com). The SPAdes plugin of Geneious Prime and the Pharokka (https://github.com/gbouras13/pharokka) version 1.6.1 were used to perform assembly and annotation respectively ^65^.

#### 4.3.3 Phage visualization

Phage structure was visualized by Transmission Electron Microscopy (TEM) as described before^19^. Briefly, 1 ml of 10^8^ PFU was centrifuged at 20,000 g (centrifuge 5430 R, rotor FA- 45-24-11HS; Eppendorf, Hamburg, Germany) for 2h at room temperature. The pellet was resuspended in 200 μl of 5 mM MgSO_4_, spotted on a 2% uranyl acetate carbon- coated copper grid and incubated for 1 min. Phage visualization was carried out with a TEM 1400 plus (Joel, Tokyo, Japan) and images were captured with a charge-coupled device camera (Gatan Orius 600).

### 4.4 PBL

The light source was designed in-house to allow a uniform irradiation over a 96-well plate inside a humid incubator (Figure 2). The irradiating surface was composed of 460nm blue LED strips (Shenzen Dengsum Optoelectronics Co. Ltd, Shenzen, China) with High Lumens 2835 SMD chips (Epistar corp., Hsinchu, Taiwan) in a density of 120 LEDs/meter. The LEDs wavelength was further measured to be 457nm (FWHM=15) by an authorized laboratory (QCC Hazorea Calibration Technologies, Kibbutz Hazorea, Israel). The device was powered with an LED driver (LCM-40, Mean Well Enterprises CO Ltd, New Taipei City, Taiwan). Pulses, with frequency of 33KHz in 50% duty cycle were achieved with an electronic circuit based on a 555 pulse generator module. Waveform and frequency were measured and visualized with an oscilloscope (Tektronix TBS1064, Beaverton, OR) connected to a light detector. The aluminum plate on which the LED strips were mounted served also to dissipate excess heat. Light intensity was measured in mW/cm^2^ at the distance of the 96-well plate with a LaserMate power-meter (Coherent, Auburn group, Coherent-Europe, Utrecht, Holand). The PBL frequency and initial power density were based on published experiments by Enwemeka et al in collaboration with Carewear©^31,33,34^.

### 4.5 Quantification Methods

#### 4.5.1 CFU

Samples were serially diluted 10-fold on LB agar plates and grown overnight in 37°C. The number of colonies was counted, and the concentration of CFU was calculated.

#### 4.5.2 PFU

The lysate was diluted 10-fold in LB broth and spotted over a PA14 bacterial lawn grown on soft 0.6% agar, and incubated overnight at 37°C. Observed plaques were counted, and the concentration of PFU was calculated.

#### 4.5.3 Growth curves

Lytic activity was assessed by inoculating logarithmic PA14 (10^7^ CFU/ml) with the experimental samples and measuring turbidity with a plate reader (Synergy; BioTek, Winooski, VT) at OD_600nm_ at 37°C with 5-second shaking every 20 min for 24 hours. The data was smoothed by a 4-point rolling average. The threshold for growth curve detection was based on the measurements of the control group. The ‘lag time’, defined as the time in hours in which the OD increased over the detection threshold, and the area under the curve at 20 hours (‘AUC 20h’) that represents the growth potential (Figure 1B) were automatically extracted using the python code ‘growth_curve_outcomes‘ that we have developed for this study and can be easily applied to plate reader growth curve data. The code can be freely accessed in github (https://github.com/bioimagehuji/growth_curve_outcomes/).

#### 4.5.4 Biofilm biomass detection

Biofilm biomass was measured using crystal violet stain ^66^ (Glentham life sciences, Corsham, UK). The liquid containing unattached bacteria was removed by pipetting, wells were rinsed with 100 μl 0.9% NaCl, and dried. A solution of 0.1% crystal violet was added to the wells (125 μl per well), incubated for 15 minutes at room temperature. Dye was removed and the wells were rinsed twice in a distilled water bath. The plate was dried at 37 °C for 1 hour and color was extracted with 30% acetic acid (100 μl per well). The purple colored solution was transferred into a new plate and OD_550nm_ was determined in a plate reader (Synergy)^66^.

#### 4.5.5 Live-dead stain

Biofilm viability was measured using the Live/Dead cell viability kit (Invitrogen, Waltham, MA) according to the manufacturer’s instructions. Propidium iodide stained the dead cells in red while SYTO9 stained the live cells in green (measured at 630 and 520 nm respectively). The fluorescence emissions of the samples were detected by a Zeiss LSM 410 confocal laser microscope (Carl Zeiss, Oberkochen, Germany) at X4 and X20 magnification. Maximal intensity of the Z-stack of each color was established from multiple horizontal optical sections (17 sections at 12.5-μm intervals for X4 and 9 sections at 10 μm intervals for X20). Three separate fields per sample were measured using ImageJ and the percentage of live bacteria of the total were calculated according to Robertson et al^67^.

#### 4.5.6 Statistics

Data is presented as mean±SD or median[interquartile rage] as appropriate. The results were the mean of five independent experiments, each an average of triplicates (unless otherwise stated). The direct effect of PBL over phages was tested by 1-way ANOVA. The role of the time of phage addition or phage concentration in combination with PBL was tested with 2- way ANOVA. The role of order of interventions, dose, and contribution of phage to the bactericidal effect over biofilm biomass was tested by 3-way ANOVA. Tukey’s Honestly-Significant-Difference Test was used for post-hoc pairwise comparisons. The role of the interventions over %Live (ratio of the number of live bacteria out of the total live+dead) was tested by Kruskal-Wallis with Conover-Inman as post-hoc. *p-value* < 0.05 was considered significant. Statistical analyses were performed with SYSTAT, version 13 (Systat Software, Chicago, USA).

## List of Supplementary Materials

- Supplemental Table 1: PBL effect on phage stability
- Supplemental Table 2: The effect of timing of phage addition
- Supplemental Table 3: The effect of combinatorial treatment on antibiotic resistance bacteria PATZ2
- Supplemental Table 4: The effect of PBL and phage on biofilm viability

## Acknowledgments

The authors gratefully acknowledge the expert technical assistance of Eduard Berenshtein from the Electron Microscopy Laboratory, Abed Nasereddin and Idit Shiff from the Genomics Applications Laboratory, and Yariv Marom of The Core Research Facility (CRF), Faculty of Medicine.

## Author contributions

All authors made significant contributions to the preparation of this manuscript. AR, RH, and LG conceptualized and drafted the manuscript, overseeing the funding acquisition process. SCG have edited the manuscript and offered valuable conceptual advice. JB, OY, SAO, and AS were instrumental in executing various methodologies, collaborating closely with AR and LG. Additionally, AR, YE, and LG were responsible in visualizing the data. Supervision throughout the project was provided by RH and LG

## Data and materials availability

Phage genomes were deposited in the https://www.ncbi.nlm.nih.gov/genbank/, accession numbers are MT933737.1 (PASA16) and PP067092 (PAShipCat1). The datasets generated during and/or analyzed during the current study are available from the corresponding author upon reasonable request.

The growth_curve_outcomes python tool can be accessed in github https://github.com/bioimagehuji/growth_curve_outcomes/

## References and Notes

1. Murray, C. J. L., et al. Global burden of bacterial antimicrobial resistance in 2019: a systematic analysis. Lancet (2022).

2. Galán, J.-C., González-Candelas, F., Rolain, J.-M. & Cantón, R. Antibiotics as selectors and accelerators of diversity in the mechanisms of resistance: from the resistome to genetic plasticity in the β-lactamases world. Front. Microbiol. 4, 9 (2013).

3. Heilig, S., Lee, P. & Breslow, L. Curtailing antibiotic use in agriculture: it is time for action: this use contributes to bacterial resistance in humans. West. J. Med. 176, 9 (2002).

4. Schuts, E. C., et al. Current evidence on hospital antimicrobial stewardship objectives: a systematic review and meta-analysis. Lancet Infect. Dis. 16, 847–856 (2016).

5. Ciofu, O., Rojo-Molinero, E., Macià, M. D. & Oliver, A. Antibiotic treatment of biofilm infections. Apmis 125, 304–319 (2017).

6. Algburi, A., Comito, N., Kashtanov, D., Dicks, L. M. T. & Chikindas, M. L. Control of biofilm formation: antibiotics and beyond. Appl. Environ. Microbiol. 83, e02508–16 (2017).

7. Pouget, C., et al. Biofilms in diabetic foot ulcers: significance and clinical relevance. Microorganisms 8, 1580 (2020).

8. Costerton, J. W., Montanaro, L. & Arciola, C. R. Biofilm in implant infections: its production and regulation. Int. J. Artif. Organs 28, 1062–1068 (2005).

9. Flemming, H.-C. & Wingender, J. The biofilm matrix. Nat. Rev. Microbiol. 8, 623– 633 (2010).

10. Mulcahy, L. R., Isabella, V. M. & Lewis, K. Pseudomonas aeruginosa biofilms in disease. Microb. Ecol. 68, 1–12 (2014).

11. Chanishvili, N. Phage therapy—history from Twort and d’Herelle through Soviet experience to current approaches. in Advances in virus research vol. 83 3–40 (Elsevier, 2012).

12. Strathdee, S. A., Hatfull, G. F., Mutalik, V. K. & Schooley, R. T. Phage therapy: From biological mechanisms to future directions. Cell 186, 17–31 (2023).

13. Pires, D. P., Meneses, L., Brandão, A. C. & Azeredo, J. An overview of the current state of phage therapy for the treatment of biofilm-related infections. Curr. Opin. Virol. 53, 101209 (2022).

14. Meile, S., Du, J., Dunne, M., Kilcher, S. & Loessner, M. J. Engineering therapeutic phages for enhanced antibacterial efficacy. Curr. Opin. Virol. 52, 182–191 (2022).

15. Onallah, H., et al. Refractory Pseudomonas aeruginosa infections treated with phage PASA16: A compassionate use case series. Med 4, 600–611 (2023).

16. Pires, D. P., Vilas Boas, D., Sillankorva, S. & Azeredo, J. Phage therapy: a step forward in the treatment of Pseudomonas aeruginosa infections. J. Virol. 89, 7449– 7456 (2015).

17. Technophage, S. Bacteriophage Therapy TP-102 in Diabetic Foot Ulcers (REVERSE). https://clinicaltrials.gov/ct2/show/NCT04803708 (2021).

18. Azam, A. H. & Tanji, Y. Peculiarities of Staphylococcus aureus phages and their possible application in phage therapy. Appl. Microbiol. Biotechnol. 103, 4279–4289 (2019).

19. Rimon, A., et al. Topical phage therapy in a mouse model of Cutibacterium acnes-induced acne-like lesions. Nat. Commun. 14, 1005 (2023).

20. Kim, M. J., Eun, D. H., Kim, S. M., Kim, J. & Lee, W. J. Efficacy of bacteriophages in Propionibacterium acnes-induced inflammation in mice. Ann. Dermatol. 31, 22–28 (2019).

21. Liu, Y., et al. Application of bacteriophage φPaP11-13 attenuates rat Cutibacterium acnes infection lesions by promoting keratinocytes apoptosis via inhibiting PI3K/Akt pathway. Microbiol. Spectr. e02838–23 (2024).

22. McCallin, S., Sacher, J. C., Zheng, J. & Chan, B. K. Current state of compassionate phage therapy. Viruses 11, 343 (2019).

23. Onallah, H., Hazan, R. & Nir-Paz, R. Compassionate Use of Bacteriophages for Failed Persistent Infections During the First 5 Years of the Israeli Phage Therapy Center. in Open Forum Infectious Diseases vol. 10 ofad221 (Oxford University Press US, 2023).

24. Young, M. J., et al. Phage therapy for diabetic foot infection: A case series. Clin. Ther. 45, 797–801 (2023).

25. Djebara, S., et al. Processing phage therapy requests in a Brussels military hospital: Lessons identified. Viruses 11, 265 (2019).

26. Yerushalmy, O., et al. Towards Standardization of Phage Susceptibility Testing: The Israeli Phage Therapy Center “Clinical Phage Microbiology”—A Pipeline Proposal. Clin. Infect. Dis. 77, S337–S351 (2023).

27. Nir-Paz, R., et al. Successful Treatment of Antibiotic-resistant, Poly-microbial Bone Infection With Bacteriophages and Antibiotics Combination. Clin. Infect. Dis. 69, 2015–2018 (2019).

28. Wang, Y., et al. Antimicrobial blue light inactivation of gram-negative pathogens in biofilms: in vitro and in vivo studies. J. Infect. Dis. 213, 1380–1387 (2016).

29. Tomb, R. M., et al. Review of the comparative susceptibility of microbial species to photoinactivation using 380–480 nm violet-blue light. Photochem. Photobiol. 94, 445–458 (2018).

30. Martegani, E., Bolognese, F., Trivellin, N. & Orlandi, V. T. Effect of blue light at 410 and 455 nm on Pseudomonas aeruginosa biofilm. J. Photochem. Photobiol. B Biol. 204, 111790 (2020).

31. Bowman, C., Bumah, V. V, Niesman, I. R., Cortez, P. & Enwemeka, C. S. Structural membrane changes induced by pulsed blue light on methicillin-resistant Staphylococcus aureus (MRSA). J. Photochem. Photobiol. B Biol. 216, 112150 (2021).

32. Hadi, J., Wu, S. & Brightwell, G. Antimicrobial blue light versus pathogenic bacteria: mechanism, application in the food industry, hurdle technologies and potential resistance. Foods 9, 1895 (2020).

33. Masson-Meyers, D. S., Bumah, V. V., Castel, C., Castel, D. & Enwemeka, C. S. Pulsed 450 nm blue light significantly inactivates Propionibacterium acnes more than continuous wave blue light. J. Photochem. Photobiol. B Biol. 202, 111719 (2020).

34. Bumah, V. V., Masson-Meyers, D. S. & Enwemeka, C. S. Pulsed 450 nm blue light suppresses MRSA and Propionibacterium acnes in planktonic cultures and bacterial biofilms. J. Photochem. Photobiol. B Biol. 202, 111702 (2020).

35. Kwon, J. H. & Powderly, W. G. The post-antibiotic era is here. Science (80-.). 373, 471 (2021).

36. Alkalay-Oren, S., et al. Complete Genome Sequence of Pseudomonas aeruginosa Bacteriophage PASA16, Used in Multiple Phage Therapy Treatments Globally. Microbiol. Resour. Announc. e00092–22 (2022).

37. Canchaya, C., Proux, C., Fournous, G., Bruttin, A. & Bruessow, H. Prophage genomics. Microbiol. Mol. Biol. Rev. 67, 238–276 (2003).

38. Sheffer-Levi, S., et al. Antibiotic Susceptibility of Cutibacterium acnes Strains Isolated from Israeli Acne Patients. Acta Derm. Venereol. 100, adv00295–adv00295 (2020).

39. Mahmoudi, H., et al. Biofilm formation and antibiotic resistance in meticillin-resistant and meticillin-sensitive Staphylococcus aureus isolated from burns. J. Wound Care 28, 66–73 (2019).

40. Segall, A. M., Roach, D. R. & Strathdee, S. A. Stronger together? Perspectives on phage-antibiotic synergy in clinical applications of phage therapy. Curr. Opin. Microbiol. 51, 46–50 (2019).

41. Lin, Y., et al. Synergy of nebulized phage PEV20 and ciprofloxacin combination against Pseudomonas aeruginosa. Int. J. Pharm. 551, 158–165 (2018).

42. Knezevic, P., Curcin, S., Aleksic, V., Petrusic, M. & Vlaski, L. Phage-antibiotic synergism: a possible approach to combatting Pseudomonas aeruginosa. Res. Microbiol. 164, 55–60 (2013).

43. Chaudhry, W. N., et al. Synergy and order effects of antibiotics and phages in killing Pseudomonas aeruginosa biofilms. PLoS One 12, e0168615 (2017).

44. Phee, A., et al. Efficacy of bacteriophage treatment on Pseudomonas aeruginosa biofilms. J. Endod. 39, 364–369 (2013).

45. Kleinpenning, M. M., et al. Clinical and histological effects of blue light on normal skin. Photodermatol. Photoimmunol. Photomed. 26, 16–21 (2010).

46. Liebmann, J., Born, M. & Kolb-Bachofen, V. Blue-light irradiation regulates proliferation and differentiation in human skin cells. J. Invest. Dermatol. 130, 259–269 (2010).

47. Bumah, V. V, Masson-Meyers, D. S. & Enwemeka, C. S. Blue 470 nm light suppresses the growth of Salmonella enterica and methicillin-resistant Staphylococcus aureus (MRSA) in vitro. Lasers Surg. Med. 47, 595–601 (2015).

48. Rupel, K., et al. Blue laser light inhibits biofilm formation in vitro and in vivo by inducing oxidative stress. NPJ biofilms microbiomes 5, 29 (2019).

49. Hashmi, J. T., et al. Effect of pulsing in low-level light therapy. Lasers Surg. Med. 42, 450–466 (2010).

50. Bumah, V. V., Masson-Meyers, D. S., Tong, W., Castel, C. & Enwemeka, C. S. Optimizing the bactericidal effect of pulsed blue light on Propionibacterium acnes-a correlative fluorescence spectroscopy study. J. Photochem. Photobiol. B Biol. 202, 111701 (2020).

51. Enwemeka, C. S., Bumah, V. V & Mokili, J. L. Pulsed blue light inactivates two strains of human coronavirus. J. Photochem. Photobiol. B Biol. 222, 112282 (2021).

52. Tomb, R. M., et al. New proof-of-concept in viral inactivation: virucidal efficacy of 405 nm light against feline calicivirus as a model for norovirus decontamination. Food Environ. Virol. 9, 159–167 (2017).

53. Richardson, T. B. & Porter, C. D. Inactivation of murine leukaemia virus by exposure to visible light. Virology 341, 321–329 (2005).

54. Dhungana, G., Nepal, R., Regmi, M. & Malla, R. Pharmacokinetics and pharmacodynamics of a novel virulent Klebsiella phage Kp_Pokalde_002 in a mouse model. Front. Cell. Infect. Microbiol. 11, 684704 (2021).

55. Ryan, E. M., Gorman, S. P., Donnelly, R. F. & Gilmore, B. F. Recent advances in bacteriophage therapy: how delivery routes, formulation, concentration and timing influence the success of phage therapy. J. Pharm. Pharmacol. 63, 1253–1264 (2011).

56. Biener, G., et al. Blue/violet laser inactivates methicillin-resistant Staphylococcus aureus by altering its transmembrane potential. J. Photochem. Photobiol. B Biol. 170, 118–124 (2017).

57. Yang, P., et al. 460nm visible light irradiation eradicates MRSA via inducing prophage activation. J. Photochem. Photobiol. B Biol. 166, 311–322 (2017).

58. Cairns, B. J., Timms, A. R., Jansen, V. A. A., Connerton, I. F. & Payne, R. J. H. Quantitative models of in vitro bacteriophage–host dynamics and their application to phage therapy. PLoS Pathog. 5, e1000253 (2009).

59. Chang, R. Y. K., et al. Bacteriophage PEV20 and ciprofloxacin combination treatment enhances removal of Pseudomonas aeruginosa biofilm isolated from cystic fibrosis and wound patients. AAPS J. 21, 1–8 (2019).

60. Chegini, Z., et al. Bacteriophage therapy against Pseudomonas aeruginosa biofilms: A review. Ann. Clin. Microbiol. Antimicrob. 19, 1–17 (2020).

61. Nash, J. F. Human safety and efficacy of ultraviolet filters and sunscreen products. Dermatol. Clin. 24, 35–51 (2006).

62. Hazan, R., et al. Auto poisoning of the respiratory chain by a quorum-sensing-regulated molecule favors biofilm formation and antibiotic tolerance. Curr. Biol. 26, 195–206 (2016).

63. Müsken, M., Di Fiore, S., Römling, U. & Häussler, S. A 96-well-plate–based optical method for the quantitative and qualitative evaluation of Pseudomonas aeruginosa biofilm formation and its application to susceptibility testing. Nat. Protoc. 5, 1460– 1469 (2010).

64. Khalifa, L., et al. Targeting Enterococcus faecalis biofilms with phage therapy. Appl. Environ. Microbiol. 81, 2696–2705 (2015).

65. Bouras, G., et al. Pharokka: a fast scalable bacteriophage annotation tool. Bioinformatics 39, btac776 (2023).

66. Shteindel, N., Yankelev, D. & Gerchman, Y. High-throughput quantitative measurement of bacterial attachment kinetics on seconds time scale. Microb. Ecol. 77, 726–735 (2019).

67. Robertson, J., McGoverin, C., Vanholsbeeck, F. & Swift, S. Optimisation of the protocol for the LIVE/DEAD® BacLightTM bacterial viability kit for rapid determination of bacterial load. Front. Microbiol. 10, 801 (2019).

