## supplementary materials for "Pulsed blue light and phage therapy: A novel synergistic bactericide"

**Supplemental Tables**

**Supplemental Table 1: PBL effect on phage stability**

PASA16 phage was irradiated with PBL 7.2 mW/cm^2^ for 0, 1, 2, or 3 hours. PA14 bacteria were incubated overnight with irradiated phage. Twenty-four hours post-infection, the number of phages (PFU/ml) and the number of bacteria (CFU/ml) were counted. The summary and statistics presented in the table are a summary of 5 separate experiments (each an average of triplicate technical repeats):

| **Treatment** | **PASA16** | | **SHIPCAT1** | |
| --- | --- | --- | --- | --- |
|  | Log(CFU/ml) | Log(PFU/ml) | Log(CFU/ml) | Log(PFU/ml) |
| **No Phage** | 7.5±0.5 | - | 8.6±0.2 | - |
| **Phage** | 3.8±0.3*** | 8.5±0.2 | 7.8±0.3*** | 8.5±0.1 |
| **Ph+1h (13 J/cm^2^)** | 4±0.3*** | 8.4±0.3 | 7.8±0.3*** | 8.3±0.3 |
| **Ph+2h (26 J/cm^2^)** | 3.6±0.2*** | 8.3±0.2 | 7.7±0.2*** | 8.4±0.3 |
| **Ph+3h (39 J/cm^2^)** | 3.8±0.3*** | 8.4±0.2 | 7.6±0.3*** | 8.5±0.2 |

Data= mean ± SD; ***p<0.001 compared to no phage by 1-way ANOVA with Dunnett’s test as post hoc.; No difference in PFU levels with or without PBL (p>0.5 by 1-way ANOVA)

**Supplemental Table 2: The effect of timing of phage addition**

PAShipCat1 phage (10^6^ PFU) was added 10, 90, or 150 minutes after treatment with 4.7mW/cm^2^ (8.5 J/cm^2^) PBL and growth curves (OD_600_) were measured from which area under the curve at 20 hours was extracted. The summary and statistics presented in the table are a summary of 4 separate experiments (each an average of triplicate technical repeats):

| **Time of phage**  **addition [min]** | **Control** | **PBL (72%)*** | **Phage (49%)*,** # | **P&B (27%)*,** #,† |
| --- | --- | --- | --- | --- |
| **10** | 28.7±0.9 | 20.8±0.4 (72%) | 15.2±0.3 (53%) | 6.1±1.2 (21%) |
| **90** | 28.7±0.9 | 20.8±0.4 (72%) | 15.3±0.4 (53%) | 8.3±0.8 (29%)‡ |
| **150** | 28.7±0.9 | 20.8±0.4 (72%) | 11.9±0.3 (41%) | 9.2±0.6 (32%)‡ |

Data=mean±SD (%reduction from control); Analysis by 2-way ANOVA with Tukey as post-hoc test: Main effects (group) *, #, † p<0.001 from control, PBL, phage respectively; Interaction within P&B (time of addition) ‡p<0.01 from 10 minutes

**Supplemental Table 3: The effect of combinatorial treatment on antibiotic resistance bacteria PATZ2**

Antibiotic resistant bacteria PATZ2 were treated with either 2 μg/ml Ceftazidime or with PASA16 10⁸ PFU/ml with or without 30 minutes PBL 7.2 mW/cm^2^ and growth curves (OD_600_) were measured from which area under the curve at 20 hours was extracted. The summary and statistics presented in the table are a summary of 5 separate experiments (each an average of triplicate technical repeats):

| Group | Control | Ant | PASA16 | PBL | PBL&PASA16 |
| --- | --- | --- | --- | --- | --- |
| Median[IQR] | 23.5[6.7] | 23.4[7.9] | 9.7[1.5]* † | 14.8[2.8]* ‡ | 3.1[2.3]* †‡ |
| Mean±SD | 26.7±8 | 25.8±8.1 | 9.7±1.2* | 15.3±1.8* | 3.4±1.8*† |

p<0.05 by Kruskal-Wallis with Conovar-Inmann as post hoc or ANOVA with Tukey as post hoc: *from Control or Antibiotics; † from PBL; ‡from PASA16

**Supplemental Table 4: The effect of PBL and phage on biofilm viability**

Formed biofilms were irradiated with 26 J/cm2 PBL and live-dead fluorescence was measured 24 hours later. The %Live of total bacteria was determined and the statistics presented in the table are a summary of 5 separate experiments (each an average of triplicate technical repeats):

| Group | Control | PASA16 | PBL | PBL&PASA16 |
| --- | --- | --- | --- | --- |
| Median[IQR] | 80 [10] | 40 [10]* † | 25 [10]* † | 15 [21]* † |

*p<0.001 vs control for all; †p<0.05 vs phage alone by Kruskal-Wallis with Conover-Inman as post-hoc
